# *Drosophila* larval epidermal cells only exhibit epidermal aging when they persist to the adult stage

**DOI:** 10.1101/2020.02.21.959783

**Authors:** Yan Wang, Sirisha Burra, Michael J. Galko

## Abstract

Holometabolous insects undergo a complete transformation of the body plan from the larval to the adult stage. In *Drosophila*, this transformation includes replacement of larval epidermal cells (LECs) by adult epidermal cells (AECs). AECs in *Drosophila* undergo a rapid and stereotyped aging program where they lose both cell membranes and nuclei. Whether LEC’s are capable of undergoing aging in a manner similar to AECs remains unknown. Here, we address this question in two ways. First, we looked for hallmarks of epidermal aging in larvae that have a greatly extended third instar and/or carry mutations that would cause premature epidermal aging at the adult stage. Such larvae, irrespective of genotype, did not show any of the signs of epidermal aging observed in the adult. Second, we developed a procedure to effect a heterochronic persistence of LECs into the adult epidermal sheet. LECs embedded within the adult epidermal sheet undergo clear signs of epidermal aging; they form multinucleate cells with each other and with the surrounding AECs on the same schedule as the AECs themselves. Our data reveals that epidermal aging in holometabolous *Drosophila* is a stage-specific phenomenon and that the capacity of LECs to respond to aging signals does exist.

**Summary Statement:** We show that *Drosophila* larval epidermal cells do not age at the larval stage. They do, however, exhibit signs of aging if they persist into the adult.

## Materials and Methods

### Fly stocks and aging

*Drosophila* were reared at 25°C on standard cornmeal medium under a 12 h light-dark cycle. Aging experiments using adult flies was performed as described in Scherfer et al., 2013. Virgin females were collected and maintained on fly food for the indicated number of days.

For most of the experiments, *w*^*1118*^ was used as a control strain. Short-lived *lam*^*G262*^ mutants (Morin et al., 2001), which have accelerated epidermal aging at the adult stage, and short-lived *Atg7*^*d77*^ mutants (Juhasz et al., 2007), which have decelerated epidermal aging in adults (Scherfer et al., 2013) were used to examine aging in larval epidermis. The *A29-GAL4* larval/adult epidermal driver was isolated in a screen for Gal4 insertions expressing in the larval epidermis and was combined with *UAS-DsRed2Nuc* (Lesch et al., 2010) to label epidermal cell nuclei.

### Synthesis and use of E2M Media

The *erg2Δ::TRP1*yeast was a kind gift from Dr. Renato Paro (University of Basel, Switzerland). Culture medium with *erg2Δ::TRP1*yeast (*erg2 Medium, E2M*) was prepared using the protocol described by Katsuyama and Paro (Katsuyama and Paro, 2013). Yeast cells were harvested, heat-inactivated, and concentrated and the resulting yeast paste was mixed with an equal volume of 1.0% glucose/1.2% agar solution and, dispensed into plastic vials. Penicillin/streptomycin was added as an antibacterial agent. Flies were grown on media containing either brewer’s yeast (NM) or E2M and the embryos were transferred onto respective media. The larvae were harvested and transferred to fresh media and the epidermis of larvae grown on NM and E2M were analyzed at the indicated days.

### UV irradiation

Larvae anesthetized with ether were mounted on microscope slides so that the lateral sides were exposed to UV. The glass slides with larvae were placed in a Spectrolinker XL-1000 ultraviolet crosslinker (Spectronics Corporation). A range of UV irradiation starting from 0 (mock), 5, 15, and 20 mJ/cm^2^ (values were read using AccuMax UVC reader) at a wavelength of 254 nm was used in Figure 2C. A range of UV irradiation starting from 0 (mock), 4, 7, 11, 12, 14, 15, and 16 mJ/cm^2^ (values were read using AccuMax UVC reader) at a wavelength of 254 nm was used in Figure 3A. The UV irradiation was 15-16 mJ/cm^2^ for Figure 3C and Figure 4. The larvae after treatment were recovered on fly food. The number of pupae and eclosed adults were counted and the epidermal membranes of adults eclosed after UV treatment were analyzed.

**Figure 1.**
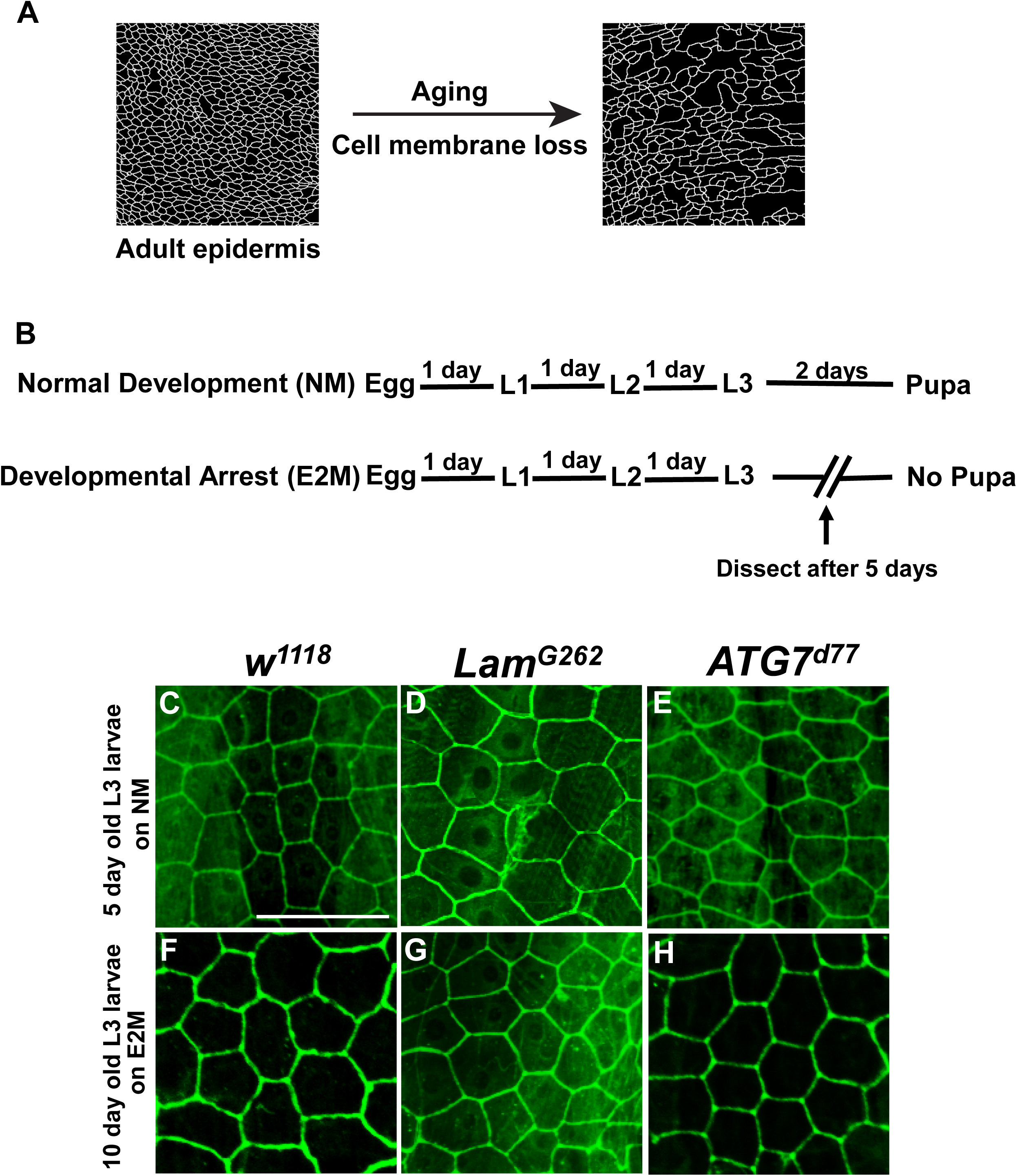
Longer-lived larvae do not exhibit signs of epidermal aging. (A) Cartoon of the morphological progression of adult epidermal aging. As the adult fly ages, white epidermal membranes are lost, leading to large blank spaces in the epidermal tissue. (B) Schematic of the timing of development on normal media (NM) and Erg2 yeast media (E2M). (C-H) Dissected epidermal whole mounts of 5-day old third instar larvae grown on NM (C-E) or E2M (F-H) and immunostained with anti-Fasciclin III to highlight epidermal cell membranes (green). *w*^*1118*^ control larvae (C, F); *lam*^*G262*^ larvae (D, G); *atg7*^*d77*^ larvae (E, H). Scale bar, 100 µm.

**Figure 2.**
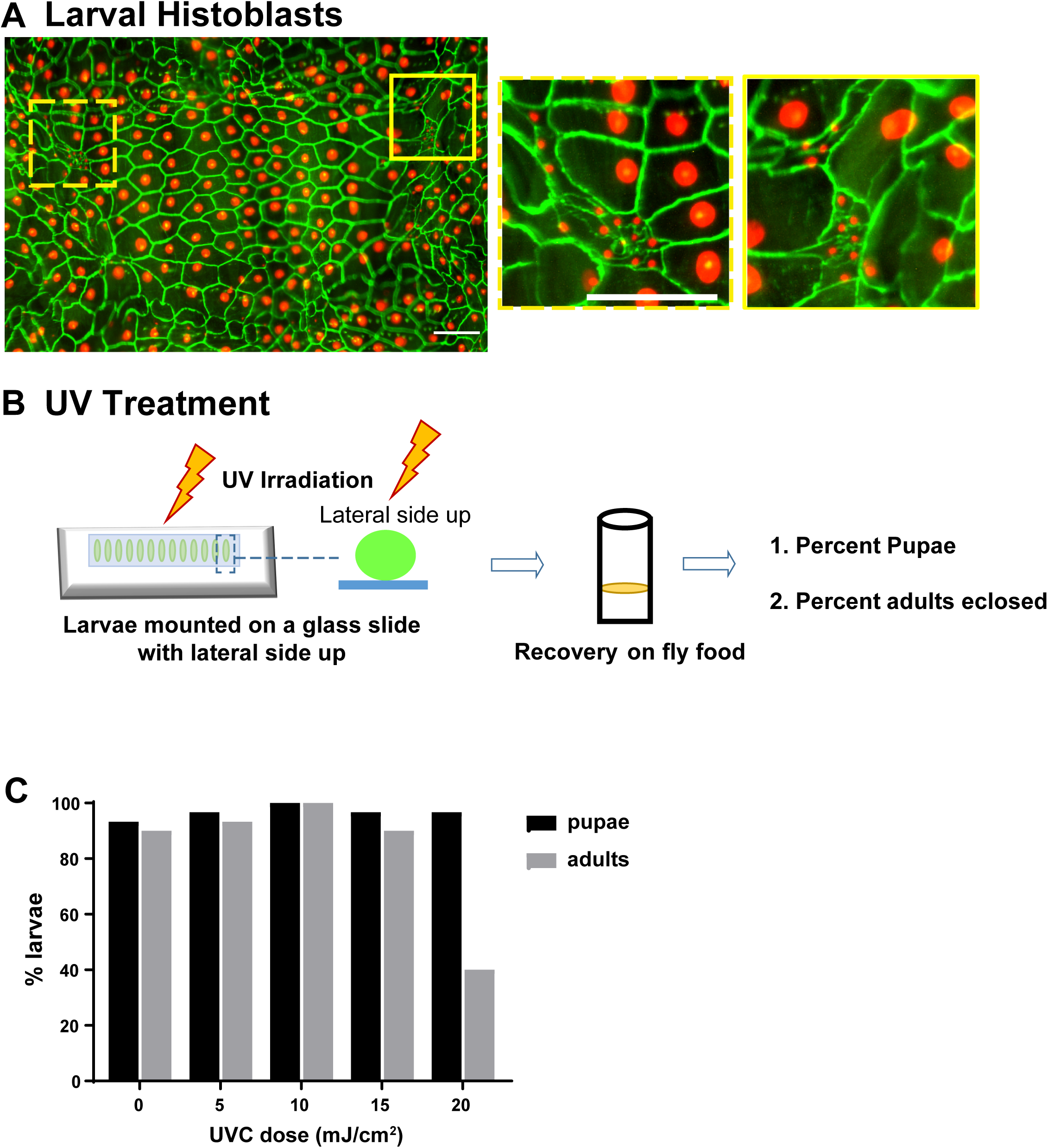
Irradiation of larval epidermal histoblasts. (A) Location and morphology of larval epidermal histoblast nests. Dissected epidermal whole mounts of third instar larvae (*A29-Gal4, UASdsRed2Nuc*) immunostained with ant-Fasciclin III to highlight epidermal membranes (green). Nuclei, red. Yellow dashed boxes highlight the epidermal histoblast nests- small clusters of cell that remain diploid within the field of larger endoreplicating larval epidermal cells (LECs). Closeups of the boxed regions show the smaller histoblast cells surrounded by their LEC neighbors. Scale bar, 100 µm. (B) Schematic of protocol for UV irradiation and recovery of larval epidermal histoblasts. (C) Quantitation of pupal (black bars) and adult (grey bars) survival following UV irradiation of third instar larvae with increasing doses (indicated) of UV. n=30.

**Figure 3.**
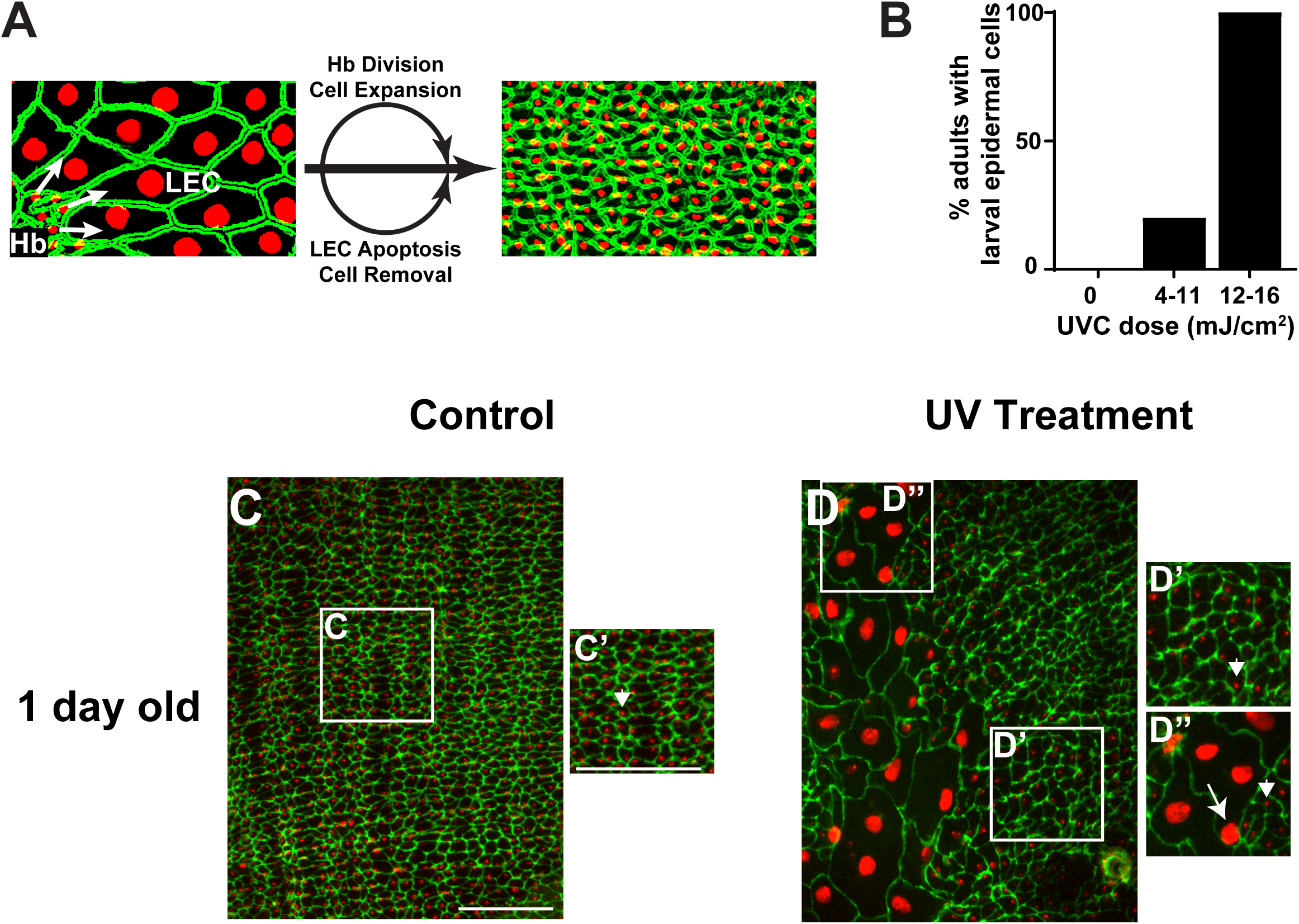
Persistence of LECs to the adult stage following histoblast irradiation. (A) Schematic of epidermal cell replacement during metamorphosis. At the pupal stage, histoblasts proliferate and migrate to replace LECs, which undergo apoptosis and are removed. (B) Quantitation of the percent of adults exhibiting large LEC-like cells as a function of the dose of UV administered during the third larval instar. (0 mJ/cm^2^: n=16; 4-11 mJ/cm^2^: n=5; 12-16 mJ/cm^2^: n=25). (C-D”) Dissected epidermal whole mounts of the one-day old adult abdominal epidermis (*A29-Gal4, UASdsRed2Nuc*) immunostained with anti-Fasciclin III (green). Nuclei, red. (C-C’) Panorama of control (non-irradiated) epidermal sheet with predominantly small mononuclear cells (before onset of epidermal aging). (C’) Closeup of white-boxed region. Arrowhead, small AEC nucleus. (D-D”) The one day old irradiated epidermis in panorama (D) exhibits both regions of exclusively small adult epidermal cells (D’) and hybrid regions that contain both large LECs and smaller AECs (D”). Arrow, large LEC nucleus. Arrowhead, small AEC nucleus. Scale bar, 100 µm.

**Figure 4.**
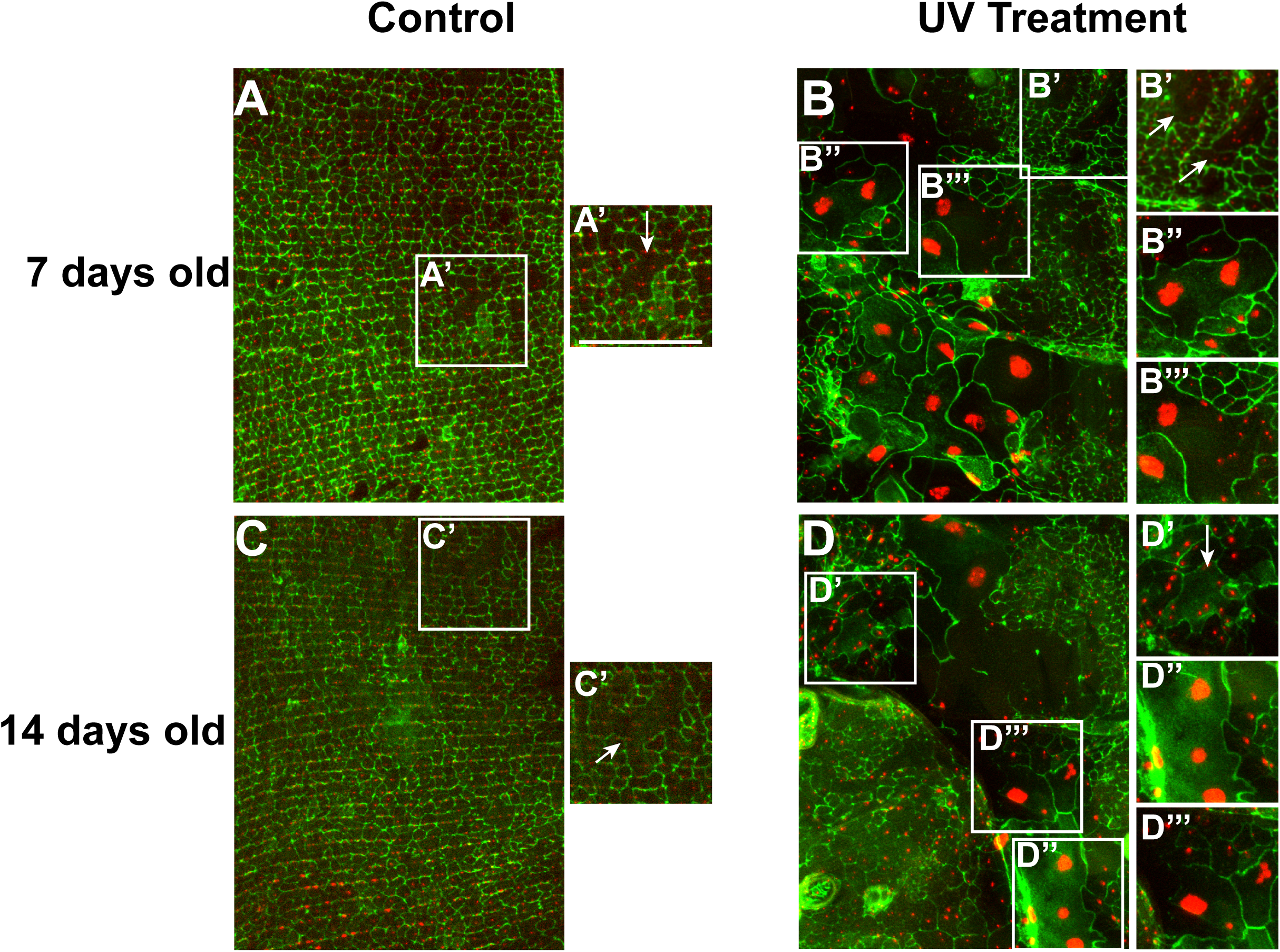
LECs embedded within the adult epidermis undergo epidermal aging. (A-D) Dissected epidermal whole mounts of seven day old adult abdominal epidermis (*A29-Gal4, UASdsRed2Nuc*) immunostained with anti-Fasciclin III (green). Nuclei, red. (A-A’) Panorama of control (non-irradiated) seven day-old epidermal sheet where epidermal aging (multinucleate epidermal cells) are now apparent (A’, arrow) Closeup of white-boxed region. (B-B’”) The seven day old irradiated epidermis in panorama (B) exhibits contains both LECs and AECs. Within the epidermal sheet there are multinucleate cells containing only small adult epidermal nuclei (B’, arrows), cells containing multiple large LEC nuclei (B”) and hybrid cells that contain both large LECs and smaller AECs (B”’). (C-C’) Panorama of control (non-irradiated) fourteen day-old epidermal sheet where epidermal aging (multinucleate epidermal cells) has progressed. (C’) Closeup of white-boxed region showing large AEC with multiple nuclei (arrows). (D-D’”) The fourteen day-old irradiated epidermis in panorama (D) exhibits contains both LECs and AECs. Within the epidermal sheet there are multinucleate cells containing only small adult epidermal nuclei (D’, arrow), cells containing multiple large LEC nuclei (D”) and hybrid cells that contain both large LECs and smaller AECs (D”’). Scale bar, 100 µm.

### Dissections, immunostaining and microscopy

Larvae were dissected and stained as previously described(Burra et al., 2013). Adult flies were anaesthetized by briefly exposing to CO_2_ and dissected as described (Scherfer et al., 2013). Briefly, flies were placed on a Sylgard plate (Dow Corning). The head and other appendices were removed using forceps and the thorax and abdomen were pinned dorsal side up using 0.1 mm diameter dissection needles (Fine Science Tools). After placing the first set of pins, 1X phosphate buffered saline (PBS) was added to the Sygalrd plate and any bubbles trapped beneath the ventral abdomen were gently flushed. Using dissecting scissors (Fine Science Tools) an incision was made and the flaps were pinned to the side. The viscera and other organs were removed and the pins were repositioned to flatten the epidermis. The dissected samples were fixed in 3.7% formaldehyde prepared in 1X PBS for 1 hour and then washed quickly with 1X PBS. After rinsing out the formaldehyde, samples were unpinned and transferred to 0.5 ml microtubes containing PHT buffer (phosphate-buffered saline containing 1% heat-inactivated normal goat serum and 0.3% Triton X100) for 1 hour before immunostaining. The samples were incubated overnight in solution containing 1: 50 dilution of mouse anti-Fasciclin III (*Drosophila* Hybridoma Bank). After primary antibody incubation, the samples were washed with PHT. Next, the samples were incubated in PHT solution containing goat anti-mouse Alexa 647 (1:500, Abcam). The samples were again washed with 1x PBS containing 0.3% Triton X100 and mounted in Vectashield mounting medium (Vector Laboratories). Larval histoblast images were captured at room temperature with a Leica MZ16 FA fluorescent stereomicroscope equipped with a PLAN APO 1.6x stereo-objective and a JENOPTIK ProgRes C14 Plus digital camera. Image Pro Plus 7.0 softwawre (Media Cybernetics) was used. Other epidermal whole mount images were captured with an Olympus FV 1000 laser scanning confocal microscope and a UPLAPO (10x/0.40 NA or 20x/0.70 NA) objective using FLUOVIEW version FV10-ASW 3.1 software at room temperature and processed with ImageJ 1.52n.

## Introduction

Holometabolous insects undergo a complete restructuring of the body plan during metamorphosis. Typically, this involves changing the morphology of a larva (usually a worm-like transitional stage) into an adult that possesses adult appendages (legs, wings, antennae) and functional reproductive organs. In some insects, including Drosophilid flies, the larval stage can enter a diapause which temporarily halts further development (Enomoto, 1981) until the local environment is conducive to further growth. In a phenomenon distinct from diapause, certain dietary restrictions that prevent synthesis of the molting hormone (Parkin and Burnet, 1986) can block the onset of pupariation and result in long-lived *Drosophila* larvae. A variety of genetic mutations also result in a greatly prolonged larval stage (Belinski-Deutsch et al., 1983, Sandoval et al., 2014), often with no pupariation or metamorphosis.

As in most insects, the *Drosophila* larval epidermis is a monolayer of large polarized polygonal epithelial cells that are adherent to an apical cuticle (Gangishetti et al., 2012) and that synthesize a basal lamina (Fessler and Fessler, 1989) separating them from the hemolymph in the open body cavity. Little is known about whether or how prolonged larval stages affect barrier tissue architecture. During metamorphosis, epidermal histoblast cells (Madhavan and Madhavan, 1980) proliferate and migrate to replace the larval epidermal cells (LECs) undergoing apoptosis (Ninov et al., 2007), thus forming a new epidermal sheet and cuticle. After eclosion, the adult epidermal cells (AECs) are substantially smaller than their larval counterparts and more rounded in shape (Scherfer et al., 2013). Like their larval counterparts (Galko and Krasnow, 2004), they are capable of undertaking physiological responses such as wound healing (Losick et al., 2013, Ramet et al., 2002). Adult flies undergo a genetically-programmed aging process in which the total average lifespan can be shortened by certain mutations (Juhasz et al., 2007, Munoz-Alarcon et al., 2007) and lengthened by others (Clancy et al., 2001, Libert et al., 2007). In a striking example of a tissue-specific aging program, many of the AECs grow thinner, lose the membranes intervening between nuclei, and eventually lose nuclei as well as adult flies age (Scherfer et al., 2013).

The question of whether larval stages can undergo a normal “aging” process, either during the normal window of development or during a prolonged version of this window, has not, to our knowledge, been addressed in the era of molecular/genetic aging research. We approached this question experimentally in two distinct ways. First, we manipulated the larval diet to create “longer-lived” larvae. We did this in control larvae, and in mutants that would normally accelerate or decelerate aging at the adult stage (Juhasz et al., 2007, Munoz-Alarcon et al., 2007). We then examined the larval barrier epidermis for the normal morphological hallmarks of adult aging (Scherfer et al., 2013)-primarily loss of membranes between intervening nuclei. Second, we developed a protocol that effects a “heterochronic” persistence of LECs into the adult epidermal sheet. We then examined whether these hybrid epidermal sheets containing both LECs and AECs, and the different cell types within them, underwent a normal process of adult skin aging. The results are presented and discussed below.

## Results

### The larval epidermis does not exhibit signs of epidermal aging

Previously, we observed that the *Drosophila* adult epidermis undergoes an age-dependent loss of cell membranes (See schematic Fig. 1A) and nuclei (Scherfer et al., 2013). To help determine whether skin-aging signals are specific to the adult stage, we asked whether larvae, a transitional juvenile form that precedes metamorphosis, also exhibit a similar age-dependent changes in the epidermis. The *Drosophila* third instar larval stage (L3) normally lasts for 2 days before the puparial molt. The time spent in this stage can be substantially prolonged through nutrient deprivation that precludes synthesis of the molting hormone (Parkin and Burnet, 1986). Larvae grown on media containing yeast mutant for the ERG2 gene cannot synthesize molting hormone and do not pupariate (Katsuyama and Paro, 2013). We developed a scheme to grow control larvae (*w*^*1118*^) and larvae mutant for aging genes on either normal media (NM) or media made with Erg2 mutant yeast (E2M) (Fig. 1B). *Drosophila* grown on normal media (NM) typically reach L3 after five days and have large polygonal epidermal cells with distinct cell membranes (Fig. 1C). On NM, both *lam*^*G262*^ mutants, which exhibit a short lifespan (Munoz-Alarcon et al., 2007) and premature/accelerated skin aging as adults (Scherfer et al., 2013) and *ATG*^*7d77*^ mutants(Juhasz et al., 2007), which exhibit decelerated skin aging as adults(Scherfer et al., 2013), exhibited a morphologically normal larval epidermis at the middle of the L3 stage (Fig. 1 D-E). When grown on E2M, ten day old L3 larvae of all genotypes tested also showed an epidermal morphology that was indistinguishable from the younger larvae grown on NM (Fig. 1 F-H). These results suggest that there is no equivalent, in larvae, to the progressive deterioration of epidermal cell membranes that is observed in the adult. This is true regardless of whether the larval genotype would or would not exhibit an aging phenotype at the adult stage.

### UV treatment of larval histoblasts to create adults with persistent larval epidermal cells

The adult epidermis is formed through proliferation and migration of larval epidermal histoblasts after the puparial molt (Madhavan and Madhavan, 1980, Ninov et al., 2007). During normal development these histoblast cells, which are diploid precursors embedded within the polyploidy larval epidermis (Fig. 2A), expand and migrate to replace dying larval epidermal cells (LECs). To interfere with this replacement process, and hopefully create *Drosophila* adults that contained persistent LECs, we developed a protocol where the lateral aspect of larvae was irradiated with UV (see methods and Fig. 2B). We first defined a UV dose that is not lethal. We observed full survival to the pupal stage up to 20 mJ/cm^2^ of UVC (Fig. 2C). Survival to the adult stage was complete up to 15 mJ/cm^2^ but dropped by over half when the dose was increased to 20 mJ/cm^2^.

Do LECs in irradiated larvae persist through metamorphosis and into the adult stage? To assess this, we irradiated larvae that carried an epidermal Gal4 driver (*A29-Gal4*) and a nuclear-localized red fluorescent transgene (*UAS-dsRed2Nuc*) (Lesch et al., 2010) that allowed us to distinguish between LECs (large cells and nuclei; 1836 ± 639 µm^2^ in cell area; n = 166) and AECs (small cells and nuclei; 138 ± 34 µm^2^ in cell area; n = 77). We hypothesize that targeted UV irradiation might disrupt the normal replacement of LECs by dividing histoblasts during metamorphosis (See schematic, Fig. 3A) In the absence of irradiation, no persistent LECs were observed (Fig. 3B) and all of the epidermal cells in the 1 day old adult epidermal monolayer had the small size and faintly-labeled nuclei characteristic of AECs (Fig. 3C-C’). At this early stage after eclosion, the loss of epidermal cell membranes has not yet commenced, multinucleate cells are rare(Scherfer et al., 2013), and no cells with the large size and characteristic large nuclei of LECs (Lesch et al., 2010, Wang et al., 2015) are apparent. At a UV dose of 4-11 mJ/cm^2^, 20 % of adults showed persistent LECs (Fig. 3B). When this dose was in the range of 12-16 mJ/cm^2^, still a dose that gives full survival to the adult stage (Fig. 2C), nearly 100 % of the adults that eclosed had persistent LECs embedded within their adult epidermis. The resulting adult epidermal sheets are shown in Fig. 3D-D”. Persistent LECs (Fig. 3D, D”, arrow) were much larger than adjacent AECs (Fig. 3D’, arrowhead), in terms of area, size, and brightness of the nucleus. Most of these cells, on day 1, were mononucleate, though some were bi- or trinucleate (Fig. 3D, D”), possibly a result of the prior UV irradiation (Babcock et al., 2009). Procedurally, the technique developed here provides the functional equivalent of a heterochronic transplant of LECs into the adult epidermal sheet, creating adults that possess a hybrid epidermis consisting of both persistent LECs and resident AECs.

### LECs that persist until the adult stage do undergo epidermal aging

We next asked what happens to these persistent LECs and the resident AECs as the adults with a hybrid epidermal sheet age. If LECs are immune to adult epidermal aging signals, as might be suggested by their behavior in longer-lived larvae, the expectation is that these cells would stay primarily mononucleate as the adult ages. On the contrary, if LECs can respond to epidermal aging signals when embedded in the adult epidermis we would expect them to lose membranes both between themselves and the surrounding AECs. The latter is what we observed. In the week-old unirradiated adult epidermis (Fig. 4A), AECs began to lose some of the membranes intervening between the cells, resulting in the previously observed skin aging phenotype-multinucleate cells (Fig. 4A’, arrow). We observed three types of multinucleate cells following irradiation in the week-old hybrid epidermis (Fig. 4B). First (Fig. 4B’, arrows), we saw the ‘standard’ adult epidermal aging phenotype-multinucleate cells consisting solely of small AEC nuclei. Second (Fig. 4B”), we saw multinucleate cells that contained two (or sometimes more) large LEC nuclei. Third, and most tellingly (Fig. 4B”), we observed multinucleate cells that contained both large LEC nuclei and small AEC nuclei (the closeup in Fig. 4B” shows a section of a cell that contains one large LEC nucleus and 9 smaller AEC nuclei, just within the highlighted region).

The same diversity of cell types is present as the adult ages to two weeks. In the absence of irradiation the loss of epidermal membranes progresses with age (compare Figure 4A, A’ to Figure 4C, C’, arrow). With irradiation, the same diverse cell hybrid cell types seen at one week (Figure 4B-B”’) are seen at two weeks (Figure 4D-D”’). The presence of these various cell types and their progression over time is strong evidence that LECs heterochronically persistent in the adult epidermal sheet can respond to epidermal aging signals with the same cellular behavior (loss of membranes between intervening nuclei) that is exhibited by AECs during epidermal aging.

## Discussion

Classical experiments in *Drosophila* and other insects have explored various aspects of whether prolonging transitional larval development stages, can impact the progression of later stages of development. For instance prolonging the larval stage in *Drosophila ampelophila* by nutrient deprivation does not lengthen the subsequent duration of metamorphosis (Northrop, 1917). Similarly, prolonging larval life in *Drosophila melanogaster* by lowering the temperature does not have a substantial effect on the duration of the subsequent adult stage (Alpatov and Pearl, 1929). In beetles (*Trogoderma glabrum*), food deprivation leads to a “retrogressive” molting where the new larvae are smaller than the previous stage. Interestingly, these larvae show physiological symptoms of senescence (increased ploidy) primarily in the fat body (Beck and Bharadwaj, 1972). Several studies have examined age-related cellular changes in the adult gut (Miquel et al., 1972), skeletal muscle(Demontis and Perrimon, 2010), and barrier epidermis (Scherfer et al., 2013), but creating adults where larval cells persist has largely been limited to examining the regenerative capacity of imaginal discs (Bryant, 1975), structures that do not persist into the adult in their larval pattern/form. Here, we sought to develop experimental paradigms that would allow us to examine “aging” in a long-lived larval stage and in larval cells heterochronically persisting into the adult stage.

Our results suggest that LECs, when held in a prolonged larval stage, do not undergo a morphological aging process comparable to that of adults. During the normal span of larval development, LECs do not show any of the hallmarks of “adult” aging-loss of membranes between nuclei or loss of nuclei themselves (Scherfer et al., 2013). Even if the larval stage is prolonged, LECs still do not show any signs of age-dependent morphological changes. One might expect *lamin* mutant larvae (Munoz-Alarcon et al., 2007), that exhibit accelerated epidermal aging in adults (Scherfer et al., 2013), to possibly show signs of epidermal aging at the larval stage. This was not observed even though the time window examined (days) is sufficient at the adult stage for AECs to have an appearance of being weeks old. LEC morphology was also unchanged in autophagy mutants (Juhasz et al., 2007) that delay epidermal aging at the adult stage (Scherfer et al., 2013) even though their adult lifespan is shortened.

Several possibilities might explain the inability of LECs to show morphological signs of aging, even when the larval stage is prolonged. One is that larvae simply do not produce the systemic signal(s) that might accompany normal adult aging. A second is that these signals exist, but that LECs are not responsive to them at this stage. Our heterochronic transplantation experiment tested this latter possibility. By developing a protocol to create adult flies that harbor persistent LECs we were able to examine the cellular morphology of LECs in a tissue that normally undergoes an age-related morphological progression. The resulting epidermis, a hybrid of LECs and AECs, underwent a normal adult epidermal aging program in which all cells participated. This suggests that LECs do have the capacity to respond to systemic aging signals present in the adult.

Most adult insect tissues are epithelial in nature and are not, as in vertebrates, replenished during adult life by resident stem cells. Each of these tissues has its own organ-specific aging program that can likely be monitored at the cellular level. For those tissues, like the barrier epidermis that have a contribution from nests of imaginal cells that are set aside at the larval stage, one can imagine using a similar irradiation strategy to interfere with replacement of the larval cells during metamorphosis. Another such tissue is the *Drosophila* tracheal system (Weaver and Krasnow, 2008). Although the organ-specific aging program of this tissue has not been examined in cellular detail, some details of the replacement process are known, including its dependence on FGF signaling (Chen and Krasnow, 2014), suggesting that either irradiation-based or genetic strategies (interfering with FGF signaling) might be viable strategies for creating heterochronic animals with a mixture of larval and adult tracheal cells. We hope that the experimental strategies outlined here will prove adaptable to other tissues, allowing an examination of how generalizable the result obtained here, with barrier epidermal cells, proves to be.

## Acknowledgements

We thank Dr. Christoph Scherfer for first examining the adult epidermis in our lab and Adriana P-H and the MDA Department of Genetics microscopy facility for microscopy assistance. Drs. Roger-Lopez-Bellido, Swathi Arur, and George Eisenhoffer read and commented on the manuscript.

